# Population-level neural correlates of flexible avoidance learning in medial prefrontal cortex

**DOI:** 10.1101/2022.12.31.522384

**Authors:** Benjamin Ehret, Roman Boehringer, Elizabeth A. Amadei, Maria R. Cervera, Christian Henning, Aniruddh Galgali, Valerio Mante, Benjamin F. Grewe

## Abstract

The medial prefrontal cortex (mPFC) has been proposed to link sensory inputs and behavioral outputs to mediate the execution of learned behaviors. However, how such a link is implemented has remained unclear. To measure prefrontal neural correlates of sensory stimuli and learned behaviors, we performed population calcium imaging during a novel tone-signaled active avoidance paradigm in mice. We developed a novel analysis approach based on dimensionality reduction and decoding that allowed us to identify and isolate population activity patterns related the tone stimulus, learned avoidance actions and general motion. While tone-related activity was not informative about behavior, avoidance-related activity was predictive of upcoming avoidance actions. Moreover, avoidance-related activity distinguished between two different learned avoidance actions, consistent with a model in which mPFC contributes to the selection between different goal-directed actions. Overall, our results suggest that mPFC circuit dynamics transform sensory inputs into specific behavioral outputs through distributed population-level computations.

## Main

Learning to appropriately respond to sensory information that is predictive of threats or rewards is a vital skill for every animal. This learning process depends on a network of interconnected brain regions involved in diverse functions such as sensory processing, the learning of stimulus-outcome associations and behavioral execution. In rodents, the medial prefrontal cortex (mPFC) has been implicated in linking sensory information to appropriate actions during learning and behavior execution in various forms of conditioning^1^. Specifically, mPFC neurons acquire strong and temporally precise responses to behaviorally-relevant stimuli over learning^1–3^. Moreover, optogenetic manipulations of prefrontal activity can drive and/or inhibit behavioral execution in a variety of paradigms such as fear conditioning^4–6^, active avoidance^7,8^, reward-based conditioning^3,9^ and conditioned place preference^10^.

While it is well established that (1) behaviorally-relevant sensory stimuli can elicit mPFC activity and (2) that such activity can influence behavior, it is still unclear how sensory-evoked mPFC activity is locally organized and transformed to drive specific actions^1-10^. Investigating the intricate nature of this transformation has been challenging due to multiple properties of mPFC neural activity and limitations of traditional experimental strategies and analysis approaches. First, learned, action-related activity is hard to distinguish from the pronounced general motion-related activity found in mPFC^11–13^. Second, stimuli and behavioral responses often show a temporal overlap inherent to task design, complicating the isolation of sensory- and behavior-related neural activity. Third, prefrontal neurons might show mixed selectivity to multiple task variables^14^. And finally, due to this temporal and spatial mixing, optogenetic approaches have limited ability to manipulate specific task-related signals as these do not necessarily align with cell types or projection-specific subpopulations that could be targeted selectively.

Here we addressed these issues by performing large-scale neuronal recordings during a mouse active avoidance paradigm with changing contingencies between stimulus and conditioned responses. This experimental approach, combined with a novel data analysis pipeline, allowed us to isolate neural correlates of individual task variables and to study changes in the neural correlates of stimuli and behaviors throughout learning.

## Results

### A novel active avoidance paradigm allows linking a sensory stimulus to two different behavioral responses

We first developed a novel 11-day instrumental conditioning paradigm for mice that we refer to as two-dimensional active avoidance. The paradigm consisted of habituation (day 1), active avoidance training (days 2-9) and extinction (days 10-11; Fig. 1A). Each session comprised 50 trials each starting with the presentation of a tone (max. duration 10 s, 80 dB, 8 kHz). In active avoidance sessions, the tone was followed by an aversive foot shock (max. duration 5 s, 0.2 mA). On each trial, mice could avoid the shock by shuttling between two compartments of the box during the tone, which immediately terminated the trial (Fig. 1B). On days 2-4, mice were required to shuttle along the x-axis of the box (Fig. 1A, Task 1, Supplementary Movie 1). To study whether and how subjects could flexibly adapt their avoidance behavior, days 5-9 required shuttling along the perpendicular y-axis (Fig. 1A, Task 2, Supplementary Movie 1). In the following text we refer to trials that were terminated by the execution of the correct shuttle action during the tone as avoid trials, and to trials that included a shock presentation as error trials (Fig. 1B). During task 1, the proportion of avoid trials increased from 40±4% to 84±2% (mean ± s.e.m., Fig. 1C). After the task switch on day 5, performance dropped to 40±6% but recovered to 81±4% by the end of task 2. This recovery was based on mice adjusting their shuttle behavior towards the correct direction (Fig. 1D, Fig. S1). While the y-shuttling rate increased from 19±6% to 81±4% between days 4 and 9, x-shuttling concurrently dropped from 84±2% to 27±4%.

**Figure 1:**
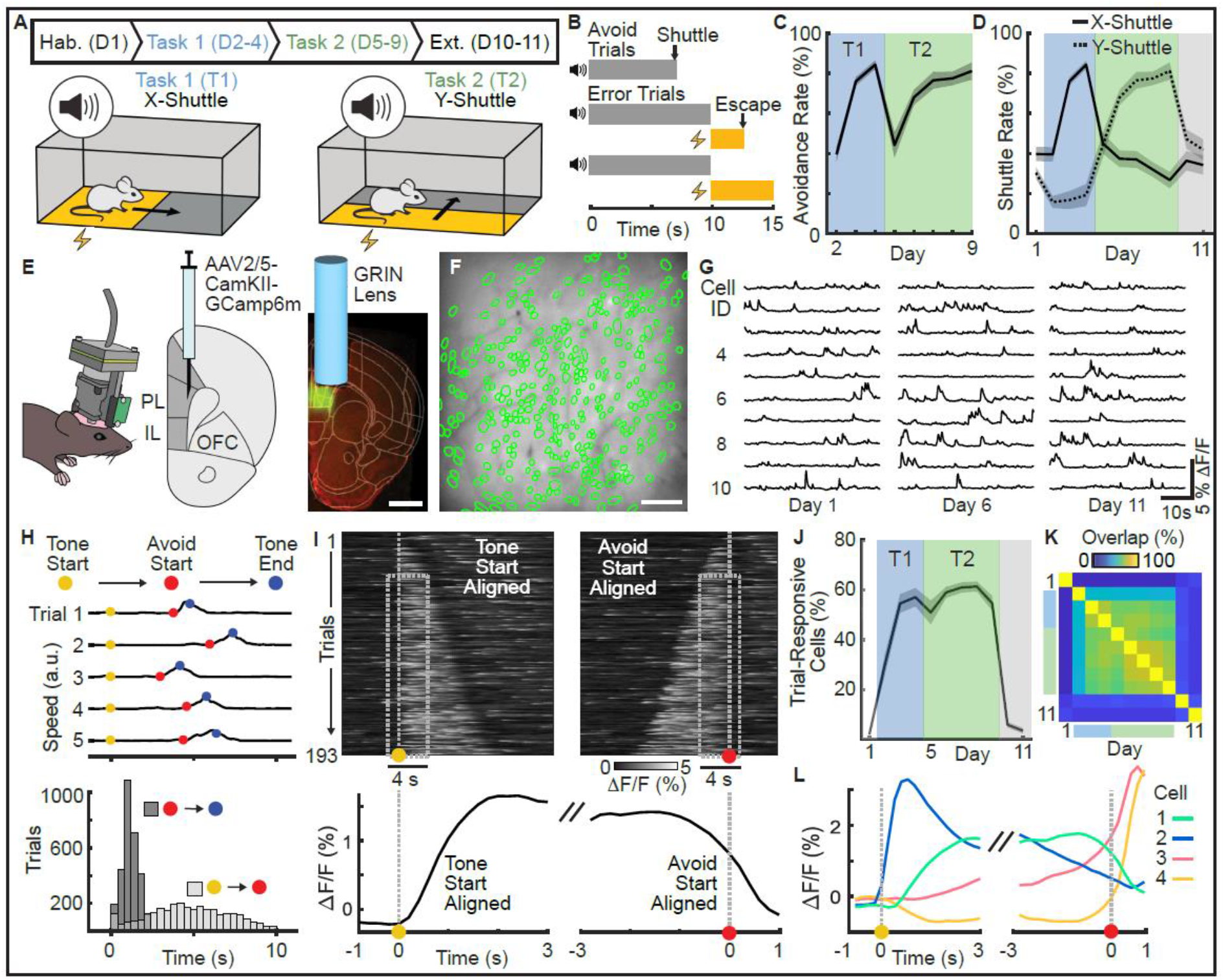
The two-dimensional Active Avoidance paradigm and recording of prefrontal population activity. (**A**) Task schematic and time course of the 11-day learning paradigm. Tasks 1 and 2 are defined by shuttling along the x and y axes of the shuttle box, respectively. (**B**) Trial structure and illustration of the different trial types (avoid and error). (**C**) Percentage of successful avoid trials per active avoidance session (n=12 mice, mean ± s.e.m.). (**D**) Shuttle rates for x-shuttles (solid line) and y-shuttles (dashed line) across 11-days of learning (n=12 mice, mean ± s.e.m.). (**E**) Miniaturized (single photon) population calcium imaging in freely behaving mice. GCaMP6m was genetically expressed in pyramidal neurons, and a gradient index (GRIN) lens was implanted above the prelimbic cortex (PL). OL, orbitofrontal cortex, IL infralimbic cortex. Scale bar: 1 mm (**F**) Cell map of an example animal. Scale bar: 100 μm. (**G**) Calcium fluorescence traces of 10 example neurons on days 1, 6 and 11. (**H**) Top: Mouse speed for five example avoid trials including markers for three reference time points (tone start, avoid start and tone end). Bottom: Distributions of latencies from tone start to avoid start and avoid start to tone end over all avoid trials (days 2-9, 12 mice). (**I**) Top: Calcium fluorescence traces of one example neuron aligned to tone start (left) or avoid start (right). Trials are sorted according to trial length. Bottom: Trial-averaged neuronal activity of the same neuron. (**J**) Percentage of trial responsive neurons across 11 days of learning (n=12 mice, mean ± s.e.m.). (**K**) Overlap of trial responsive sub-populations across 11 days. (**L**) Trial-averaged response of four example neurons aligned to tone start (left) or avoid start (right).

To investigate the neural correlates of the learned avoidance behaviors in mPFC, we expressed the genetically encoded calcium indicator GCaMP6m in excitatory neurons of the prelimbic area (Fig. 1E, Fig. S2) and used miniaturized fluorescence microscopy to image population activity in freely behaving mice (Supplementary Movie 2). This allowed us to record and track the activity of 3326 mPFC excitatory neurons of 12 mice (277 ± 49 neurons, mean ± s.d. over mice) throughout the whole 11-day paradigm (Fig. 1F, G).

To analyze the recorded neural activity during avoid trials, we first aligned recordings to two key events within each trial to account for trial-to-trial variability: tone start and avoidance action start (Fig. 1H). In a window around these alignment time points, sensory stimulation and behavior were consistent over trials, such that we could compute trial averages and jointly analyze neural responses from multiple trials (Fig. 1I). We found that during active avoidance sessions, 53±3% (mean ± s.e.m., n = 12 mice) of all recorded cells showed significantly different activity during the trial window (tone start to action start) as compared to baseline periods (Fig. 1J, mean abs. z-score > 1.96, p = 0.05, two-tailed). This fraction was substantially lower in habituation (2±1%) and extinction sessions (4±1%), and the overlap between the classified cell subsets was high between avoidance sessions (61±11%), but low between extinction sessions (14±13%, Fig. 1K). These results suggest that mPFC is recruited for sensory processing and/or production of avoidance behavior during active avoidance sessions. The responses of individual cells were highly diverse (Fig. 1L). While some cells’ activity clearly aligned to the tone or the avoidance action, other cells showed diverse temporal dynamics. Since it was difficult to isolate neuronal signals specific to the sensory stimulus, motion and avoidance action on the single cell level, we next turned to population level decoding approaches.

### Alignment of neural recordings from different mice into a joint subspace

Decoding approaches allow identifying and capturing differences in neural population activity between trial types (e.g. avoidance vs. error trials). Generally, such approaches are well suited in settings where the number of samples (here trials) exceeds the number of dimensions (here cells). In typical neuroscience settings, however, we record high-dimensional neural signals (many cells) but only have few behavioral trials per subject. To facilitate decoding analyses, we asked if we could jointly analyze trials of different subjects in a low-dimensional coding subspace that is aligned between subjects (Fig. 2A). This approach required the recorded neural activity to have two properties: (1) the high-dimensional recordings can be well described by low-dimensional trajectories in the state space spanned by the recorded cells and (2) the neural activity of subjects that perform the same task follow similar task-related dynamics. Using a dimensionality reduction and alignment procedure (details in SM), we confirmed that our data satisfies these two properties (Fig. S3). This allowed us to define a 10-dimensional joint subspace containing aligned neural trajectories for all subjects. In the following thus jointly analyze data from all subjects and perform decoding analyses in the 10-dimensional subspace.

**Figure 2:**
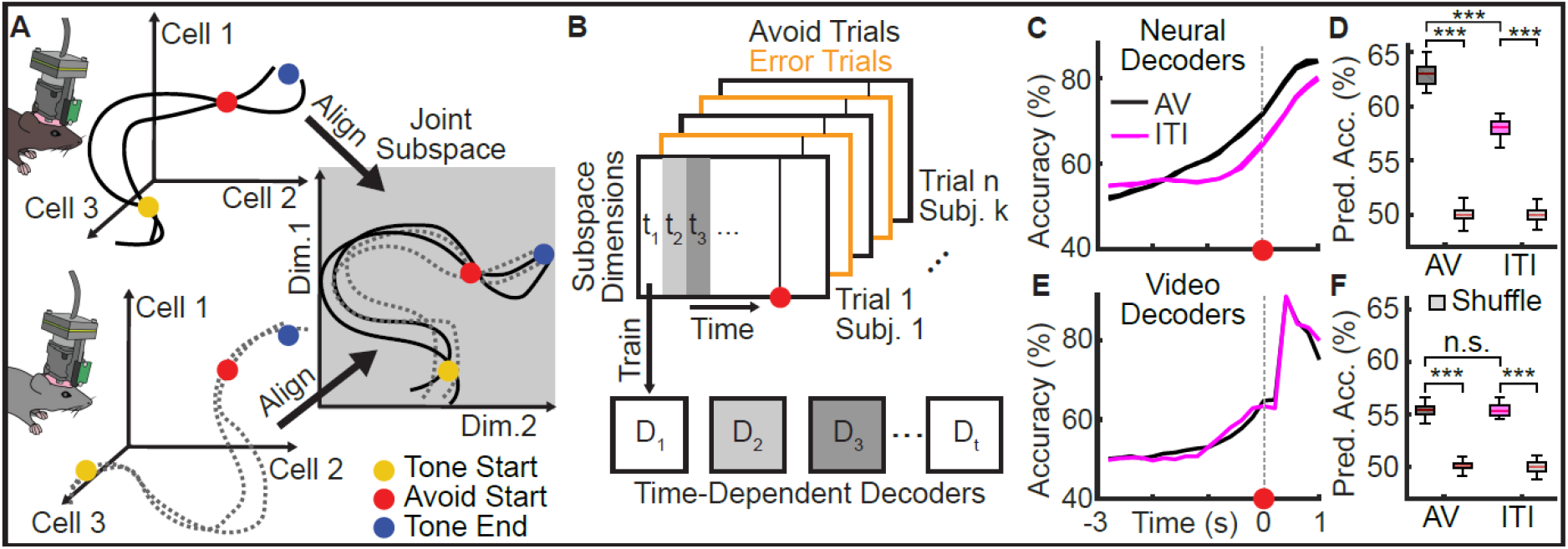
Subject alignment and prediction of avoidance actions. (**A**) Illustration of the neuronal subspace alignment procedure across animals (see Fig. S3 for details). (**B**) Schematic of the decoding approach to predict avoidance behavior from mPFC neuronal activity. For each time step an individual decoder was trained to predict the trial outcome (avoid or error). (**C**) Decoding accuracies across time for decoding of avoid vs. error trials (AV, black) and inter-trial-interval shuttles vs. random ITI periods (ITI, purple; mean ± s.e.m., n = 50 repetitions of the analysis using different samples of trials, see methods). (**D**) Quantification of predictive accuracy (time-averaged accuracy from (C) for the 2 s window before shuttle start) for the AV and ITI cases and comparison to decoding with randomly shuffled labels. (**E, F**) Same as (C) and (D), but for decoders trained using the animals’ speed extracted from video tracking data.

### Avoidance-predictive activity is distinct from activity related to general motion

To test if mPFC population activity contained predictive information about upcoming avoidance actions, we trained decoders to discriminate neural activity data from avoid and error trials projected into the joint subspace (Fig. 2B). To capture dynamical processes during the trial, we trained individual support vector machine (SVM) decoders for every time-step on temporally aligned trials. We aligned avoid trials to the action start. For error trials, however, this alignment point does not exist. We thus sampled an alignment point for each error trial, such that the distribution of trial lengths (see Fig. 1H) matched the one of avoid trials. This prevented the trial length from being informative about trial type.

Consistent with previous work^8^, we found that decoding accuracy increased towards action start (Fig. 2C) and was above chance levels in the two second window before action start (Fig. 2D, p < 0.001 Wilcoxon ranksum test), indicating that mPFC population activity contained predictive information about avoidance actions. To test if this effect was specific to avoidance actions or was rather a general property of the shuttle motion, we trained an additional set of SVM decoders to discriminate between spontaneous shuttles in the inter-trial interval (ITI) versus randomly sampled ITI periods (Fig. 2C). The resulting accuracies also exceeded chance levels (Fig. 2D, p < 0.001, Wilcoxon ranksum test), but were lower than for the avoid versus error setting (Fig. 2D, p < 0.001, Wilcoxon ranksum test). To test whether this difference could be explained by differences in motion kinematics between ITI shuttles and avoid shuttles, we trained a set of decoders using video tracking data (Fig. 2E), which showed that the predictive information contained in the animals’ motion profile was comparable between the two settings (Fig. 2F, p = 0.91, Wilcoxon ranksum test). Together these findings show that mPFC activity encodes information about upcoming avoidance actions, which cannot solely be explained by correlates of general motion. However, it remains unclear how the neural correlates of avoidance and motion relate to each other. We thus next assessed whether we could disentangle these signals during avoidance trials.

To distinguish between motion- and avoidance-related activity, we asked if the predictive performance of avoid and ITI decoders was based on different population activity patterns. We first used principal component analysis (PCA) to identify motion dimensions as the dimensions of maximal variance during ITI shuttles (Fig. S4A-C). Next, we tested how removing these motion dimensions from the joint subspace affected decoding performance in the ITI and avoid settings. We removed motion dimensions by projecting trial data from the joint subspace into the nullspace of the considered motion dimensions. We found that removing two motion dimensions led to the largest relative drop in ITI decoding accuracies (Fig. 3A, B) and that decoding accuracy was reduced both before and after action start (Fig, 3B, p < 0.001, Wilcoxon ranksum test). The decrease of predictive accuracy was substantially lower for avoid versus error decoding (Fig. 3A, B, p < 0.001, Wilcoxon ranksum test). These results show that most of the motion-related activity is contained in a low-dimensional subspace and that avoidance decoding does not depend on activity in this subspace. Thus, avoidance-related activity must be contained in different dimensions, and we next asked if we could capture these dimensions in the remaining neuronal subspace (i.e. the nullspace of the two identified motion dimensions).

**Figure 3:**
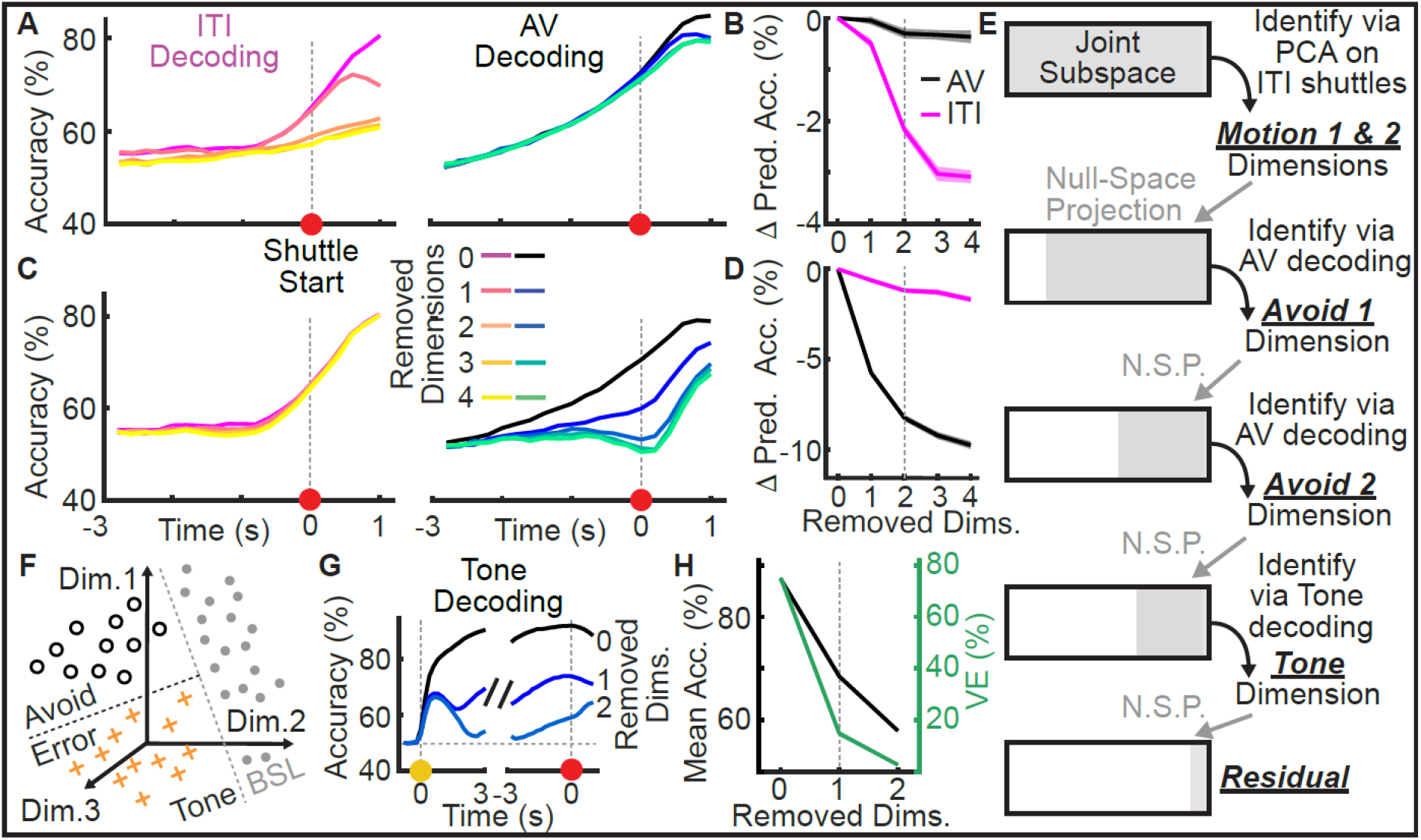
Decomposition of mPFC population activity into dimensions related to motion, avoidance actions and tone stimuli. (**A**) Mean accuracy of neural decoders for ITI (left) or avoid shuttles (right) after the progressive removal of up to four motion dimensions (n = 50 repetitions). (**B**) Drop in predictive accuracy (time-averaged accuracies from the 2 s window preceding shuttle start) of ITI and avoid decoders from A). (**C**) Mean accuracy of neural decoders for ITI (left) or avoid shuttles (right) after the progressive removal of up to four avoid dimensions (n = 50 repetitions). (**D**) Drop in predictive accuracy of ITI and avoid decoders from C). (**E**) Schematic showing the progressive decomposition of the joint subspace into five coding dimensions and a residual space. (**F**) Schematic illustrating tone vs. baseline (BSL) decoding. (**G**) Tone decoding accuracies after progressive removal of up to two tone dimensions. (**H**) Time-averaged tone decoding accuracy of the decoders from G) and variance explained for the respective decoding dimensions. In B), D) and H) lines and shaded areas correspond to mean ± s.e.m. over 50 repetitions and vertical dotted lines correspond to the number of dimensions chosen for subspace decomposition.

To identify avoidance-predictive coding dimensions, we devised an iterative approach based on decoding (Fig. S4D-F). We first projected all trial data into the motion nullspace (using 2 motion dimensions) to remove predictive information related to motion. Next, we trained a time-independent SVM decoder to discriminate between avoid and error trials and interpreted the projection axis of the decoder as an avoid dimension. To find additional avoid dimensions, we again projected trial data into the nullspace of the identified avoid dimension and repeated the process. We again evaluated the removal of the identified avoid dimensions for the ITI and avoidance settings and found that the removal of the first two avoid dimensions strongly reduced performance in the avoid versus error setting (Fig. 3C, D, p < 0.001, Wilcoxon ranksum test). In the ITI setting, predictive accuracy was also reduced (Fig. 3C, D, p < 0.001, Wilcoxon ranksum test), but the effect was substantially smaller than for the avoid versus error setting (Fig. 3C, D, p < 0.001, Wilcoxon ranksum test). Taken together, these results show that avoidance-related and motion-related activity are largely contained in orthogonal, low-dimensional subspaces.

### mPFC population activity can be decomposed into interpretable, orthogonal dimensions

In addition to avoidance and general motion, tone stimuli are a key variable during active avoidance trials. We thus asked if we could identify tone-related activity in the nullspace of the four identified motion and avoidance dimensions (Fig. 3E). We first trained SVM decoders to discriminate between tone (during avoid and error trials) and non-tone (during ITI) time periods (Fig. 3F). We found that shortly after tone onset, the decoding accuracy was consistently above 80% (Fig. 3G), indicating the presence of a reliable tone representation during the trial. To investigate the dimensionality of this tone representation we again tested the effect of iteratively removing tone decoding dimensions. Removing the first dimension decreased the mean accuracy from 87.3±0.2% to 68.8±0.3% (mean ± s.e.m., Fig. 3H). While this first dimension did not contain all tone-related information, it captured the majority (75.1±1.1%) of the remaining variance in the joint coding subspace, whereas subsequent decoding dimensions were limited to 14.9±1.1% or less. We therefore focused on this one tone dimension in subsequent analyses. Taken together, the decomposition of mPFC neuronal activity into five orthogonal dimensions (Motion 1, Motion 2, Avoid 1, Avoid 2 and Tone) constitutes a compact and interpretable representation of task-related neural activity.

To analyze how population activity in the five coding dimensions evolves over the trial, we projected the activity into each of these dimensions (Fig. 4A, top row). We found that during avoid and error trials, the activity in the two motion and the two avoid dimensions followed similar trajectories (Fig. 4B, Pearson correlation coefficient = 0.85±0.07, mean ± s.d. over 6 comparisons). Activity in these four dimensions was low at the tone start, with no differences between avoid and error trials. Activity then ramped up towards the start of the avoidance action, with a stronger increase on avoid trials compared to error trials. In contrast, activity in the tone dimension was strongly affected by tone onset and exhibited similar trajectories for avoid and error trials up to action start. Overall, the five coding dimensions captured 91±3% of the variance in avoid and error trial averages, showing that our subspace decomposition did not miss any major sources of activity (Fig. 4C). Despite the similarity of the temporal evolution of activity in the motion and avoid dimensions during the trial, there were clear differences between these dimensions for ITI shuttling (Fig. 4A, bottom row). Activity in the motion dimensions increased around the ITI shuttling start in a similar way to the avoid start. The two motion dimensions accounted for 95±1% of the variance in the population activity averaged over ITI shuttles (Fig. 4D). In contrast, the avoid dimensions only explained 2±1% of the variance, as activity was not strongly affected by ITI shuttles. These results confirm that the two identified motion dimensions indeed code for general motion, which occurs both during avoid trials and ITI, whereas the activity in the avoid dimensions is specific to avoid trials. To assess how the five coding dimensions relate to the activity of individual cells, we calculated dimension weight vectors for individual subjects by mapping the subject-specific projection matrices from the subject alignment procedure onto the five coding dimensions. Weight distributions were similar between all dimensions, were centered around zero and had long tails for both negative and positive values (Fig. 4E). We did not observe clear clustering into subpopulations and the distributions are consistent with distributed coding via mixed selectivity.

**Figure 4:**
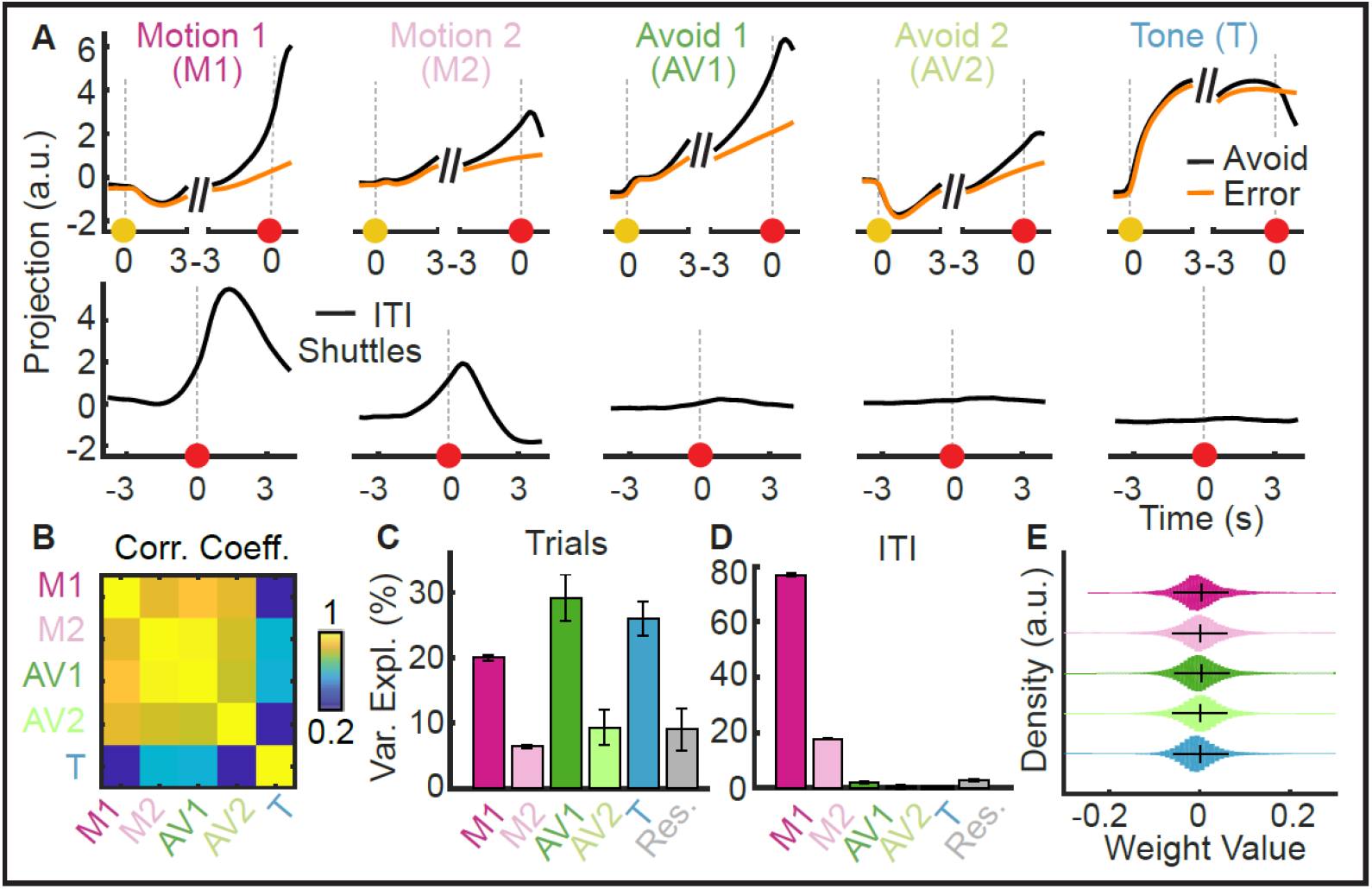
Characterization of low dimensional task-related population activity. (**A**) Mean projections (n = 50 repetitions) of neural data onto the five coding dimensions for avoid and error trials (top row) and ITI shuttles (bottom row). (**B**) Pearson correlation coefficient between pairs of coding dimension projections (avoid and error projections concatenated; mean over n=50 repetitions). (**C**) Variance explained by individual dimensions for avoid and error trials. Mean ± s.d. over 50 repetitions. (**D**) Same as C) for ITI shuttles. (**E**) Distribution of weights values that relate individual neurons to the 5 coding dimensions. Weight values are calculated per subject (see main text) and distributions were created by pooling weights over 12 subjects and 50 repetitions (Whiskers display mean and s.d.).

We next asked, how the activity in the five-dimensional coding space evolved over our learning paradigm by analyzing projections calculated for different phases of the experiment (Fig. 5A-C, Fig. S5). Motion-related activity dominated the neural variability in habituation and extinction sessions, but had a reduced relative contribution during active avoidance sessions (63.7±1.8% vs. 30.6±0.5% variance explained, Fig. 5C). In contrast, tone- and avoidance-related activity emerged in active avoidance sessions (56.1±2.8% variance explained vs. 12.0±2.2% in habituation, Fig. 5C), indicating that these activity patterns are learned and task-related. These results show that mPFC activity is engaged during active avoidance learning and develops responses to behaviorally-relevant sensory stimuli as well as activity specific to avoidance actions. Interestingly, the avoid 2 dimension seemed to be differently engaged in tasks 1 and 2, suggesting a task-related change in avoidance-related activity (Fig. 5C).

**Figure 5.**
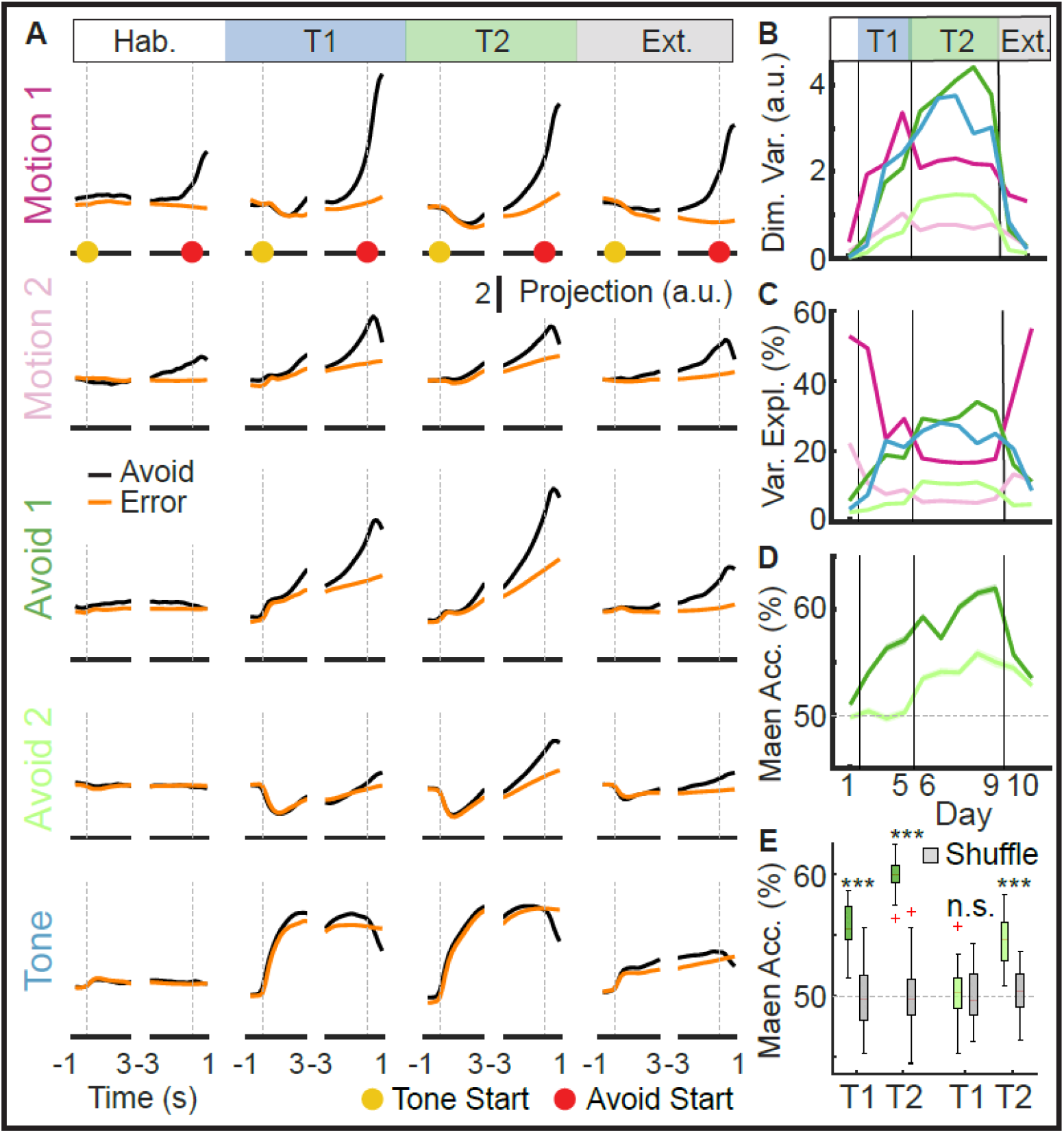
Emergence of low-dimensional, task-related neuronal signals in mPFC. (**A**) Mean projections (n = 50 repetitions) of neural data onto the five coding dimensions for avoid and error trials during habituation (day 1), task 1 (days 2-4), task 2 (days 5-9) and extinction (day 10-11). (**B**) Absolute variance within each coding dimension and (**C**) relative variance explained by the five coding dimensions across the 11-day learning paradigm. Mean ± s.e.m. over 50 repetitions. (**D**) Avoid versus error decoding accuracy for decoders evaluated on individual days. Mean ± s.e.m. over 50 repetitions. (**E**) Accuracies from (D) averaged for tasks 1 and 2 compared to accuracies obtained from decoding with shuffled data.

### Avoidance-related activity distinguishes between tasks

Based on this observation we further investigated the avoid versus error decoders that we initially used to define the avoid 1 and avoid 2 dimensions (Fig. 3, Fig. S4). To assess time-dependent changes in the decoders’ ability to discriminate avoid and error trials, we trained the decoders using data from all avoidance sessions, but tested them using data split into individual sessions (Fig. 5D). We found that the avoid 1 decoder worked best in task 2 sessions, but also showed above chance performance in task 1 (Fig. 5E, p < 0.001, Wilcoxon ranksum test). In contrast, the avoid 2 decoder performed above chance level in task 2 (p < 0.001, Wilcoxon ranksum test), but not in task 1 (Fig. 5E, p = 0.66, Wilcoxon ranksum test). This difference in decoding performance indicates that avoid 1 activity generalizes to both avoidance behaviors, while avoid 2 emerges with the task switch to accommodate the altered avoidance behavior in task 2. Taken together, these results suggest that the task switch changes mPFC coding of the avoidance action by layering additional avoidance-related activity.

To test whether the task-related changes were specific to the avoid 2 dimension or also affected other dimensions, we explicitly tested for task-based differences using an additional decoding analysis. We first trained decoders to discriminate trial data from task 1 and task 2 (task decoding, Fig. 6A), analogously to avoid versus error decoding. We trained independent sets of time-dependent task decoders for avoid trials, error trials and ITI shuttles based on the activity in the 5-dimensional coding space and found that task decoding accuracy differed between the three settings (Fig. 6B, C). Task decoding was more accurate for avoid trials than for error trials (Fig. 6C, p < 0.001, Wilcoxon ranksum test) or ITI shuttles (p < 0.001, Wilcoxon ranksum test). During avoid trials, decoding accuracy ramped up towards avoidance actions (Fig. 6B). These dynamics were less pronounced in error trials, indicating that activity informative about task identity is mostly related to the execution of avoidance actions. Although task decoding accuracy for ITI shuttles also increased towards the shuttle action, the performance was generally lower than for the avoid setting. This suggests that task decoding in avoid trials was predominantly based on avoidance-related, rather than motion-related activity.

**Figure 6:**
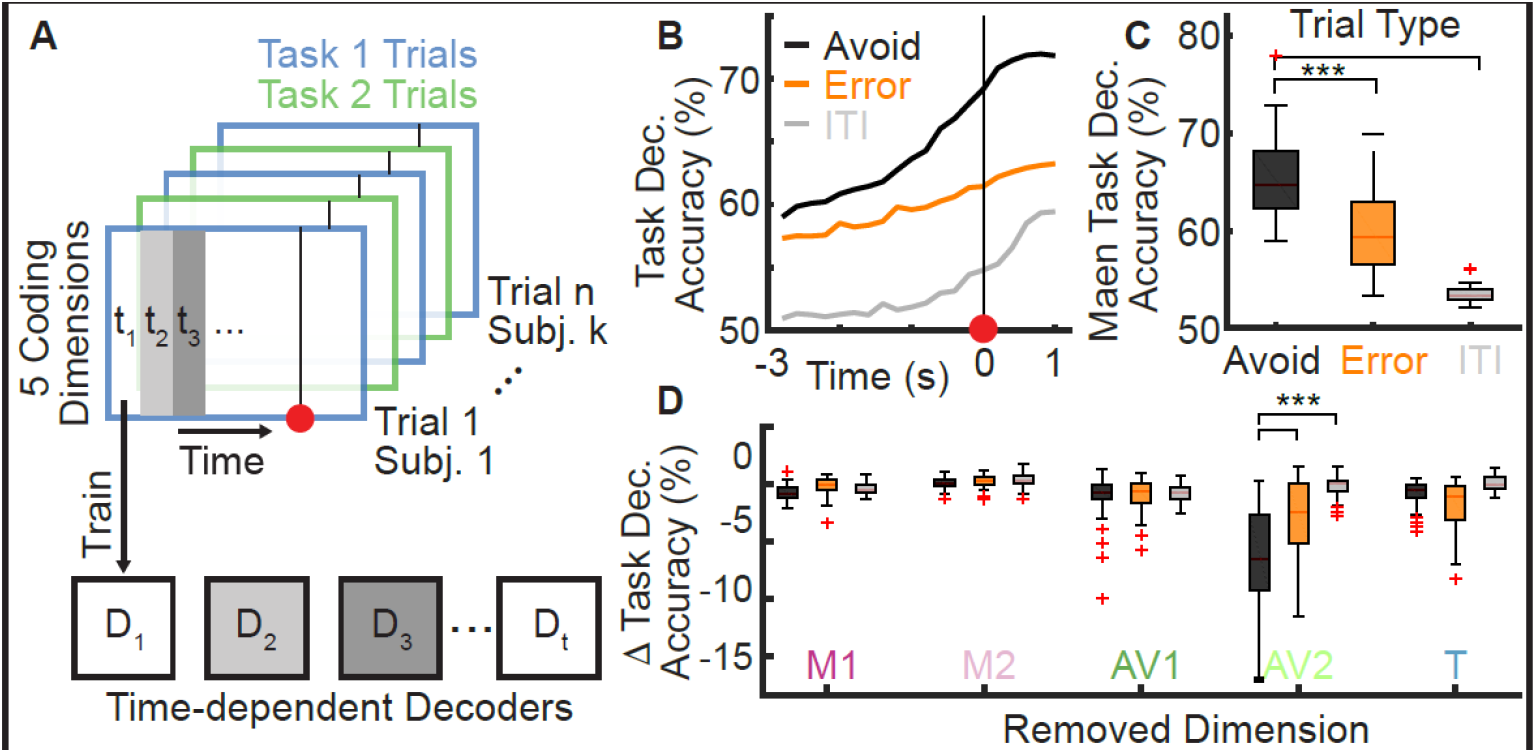
The task switch affects avoidance population coding. (**A**) Schematic of task decoding, where we trained decoders to distinguish data from tasks 1 and 2 trials. (**B**) Task decoding accuracy for time-dependent decoders trained to discriminate between task 1 and task 2 data separately for avoid trials, error trials or ITI shuttles (mean over 50 repetitions). (**C**) Temporal average for data from B). (**D**) Drop in mean decoding accuracy when removing individual coding dimensions via nullspace projection for the three task decoding settings (avoid, error and ITI; n = 50 repetitions).

To further investigate how decoding performance depended on activity in the five coding dimensions, we next asked how individually removing the respective dimensions affected task decoding accuracies. We evaluated task decoding accuracies on avoid trials, error trials and ITI shuttles. Over all dimensions and trial types, removing the avoid 2 dimension on avoid trials had the largest impact, decreasing the task decoding accuracy by 6.7±4.1% (Fig. 6D). Removing the avoid 2 dimension also caused the largest drop in accuracy for error trials (-3.1±3.6%). However, the effect was substantially smaller than for avoid trials (p < 0.001, Wilcoxon ranksum test). These results show that task-related changes were most prevalent during avoidance trials and were predominantly based on avoidance-related activity, suggesting that mPFC adjusts to changes in avoidance behavior by specifically changing its encoding within avoidance-related dimensions.

### mPFC sensory responses are modulated by avoidance behavior

Our subspace decomposition analysis shows that tone-related and avoidance-predictive activity can be decomposed into independent dimensions. Yet, we also observed that the activity in the tone dimension was modulated by the execution of avoidance actions (Fig. 7). In general, tone dimension activity was well correlated with the binary tone on/off timing for individual subjects (Fig. 7A, Pearson correlation coefficient = 0.59±0.01, calculated for individual subjects during active avoidance sessions, mean ± s.d. over 50 repetitions). However, we observed an exception at the time of action start, where the tone signal dropped 2 time steps (400 ms) after the action start (Fig. 7B), even though the tone only turned off approximately 1 s after the action start, when the action was completed (see Fig. 1H). Alignment to the end of the tone showed that the drop of activity in the tone dimension occurred 3 time steps (600 ms) prior to the actual offset of the tone (Fig. 7C). To further examine the interaction between the tone dimension and the execution of the tone-induced shuttle behavior, we next focused on a particular trial set from the transition period between tasks 1 and 2. In early task 2 trials, mice performed x-shuttles as learned in task 1, which however did not lead to avoidance anymore in task 2. During these trials, we observed a similar drop in the tone dimension activity aligned to action onset, despite the continued tone presentation (Fig. 7D, E). At 1.2 s after action onset, the tone dimension activity was decreased by 43±7% as compared to trials without shuttle actions. These results suggest that the mPFC tone representation is modulated by the execution of the learned behavior that has been associated with the termination of the tone and the avoidance of the shock.

**Figure 7:**
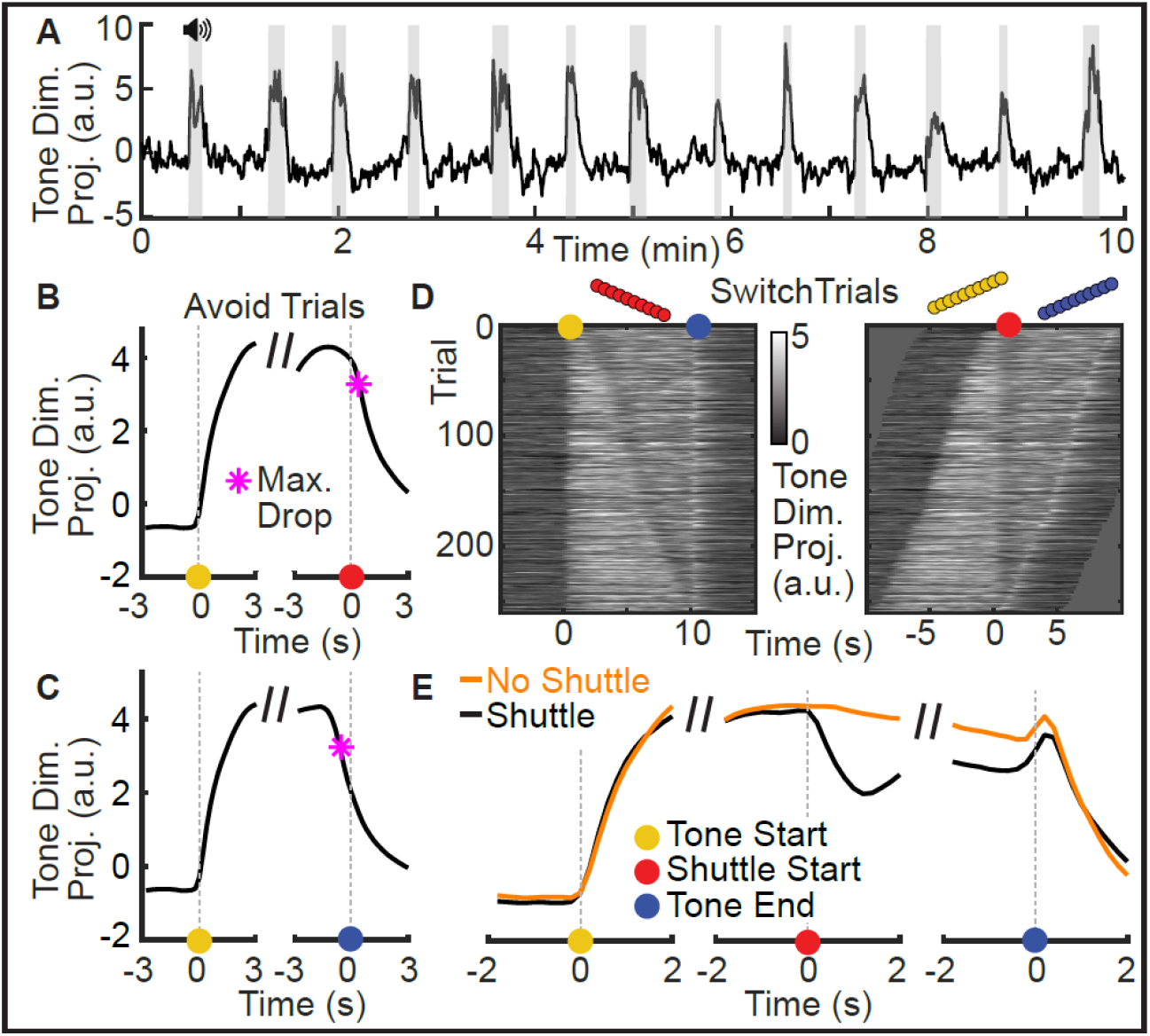
Avoidance behavior affects mPFC tone encoding. (**A**) Tone dimension projection over a 10 min time window from an example session of one mouse. Tone presentations are marked in gray. (**B**) Mean projection (n = 50 repetitions) of neural data onto the tone dimension during avoid trials aligned to tone start (left) and avoid start (right). The max. drop point (magenta star) refers to the time step prior to the maximum decrease of tone dimension activity between two consecutive time steps (5 Hz). (**C**) Same as B) but aligned to tone start (left) and tone end (right). (**D**) Tone dimension activity for switch trials (error trials from days 5 and 6 that included an x-shuttle, but no y-shuttle action). Trials were either aligned to the tone start (left) or avoid start (right) (**E**) Trial averages for data from D) (black line) aligned to tone start (left), avoid start (middle) and tone end (right). The orange line represents trials without shuttles. For these trials, the shuttle start point was randomly sampled to match the distribution of shuttle starts from the shuttle trials.

## Discussion

In this work, we developed a new two-dimensional active avoidance paradigm and combined it with large scale neural recordings in mouse mPFC and a novel data analysis approach. This allowed us to identify and characterize mPFC neural correlates of sensory stimuli and avoidance actions and to study them over learning. We show that the recorded high-dimensional population activity can be decomposed into five interpretable dimensions encoding motion, the tone and avoidance. Importantly, our approach allowed us to distinguish between learned avoidance-related activity and activity related to general motion. We show that these signals are based on orthogonal population patterns that exhibit similar dynamics during active avoidance trials but behave differently during the inter-trial interval. In addition, we found that activity in tone and avoidance dimensions emerges with learning and disappears again in extinction sessions, consistent with a model in which mPFC uses sensory driven responses to drive behavior execution. Moreover, one of the identified avoidance dimensions discriminated between the two avoidance tasks and only emerged in the second task. This suggests that the mPFC represents behaviors with sufficient resolution to enable linking stimuli to specific behavioral responses. Interestingly, we found that the execution of avoidance behaviors suppressed sensory-related activity, suggesting that mPFC sensory representations also depend on the behavior of the animal. Overall, these results point towards the mPFC implementing the sensory-behavior link through dynamically interacting neural correlates that represent essential task features and are contained within a low-dimensional subspace of the overall population activity.

The interpretation of neural activity during active avoidance trials is challenging due to the temporal overlap of sensory stimuli, cognitive processes and motor signals as well as mixed selectivity to these signals. We addressed these challenges by combining several data analysis steps that allowed us to identify and isolate distinct and well-defined neural correlates at the population level. First, we used a procedure to align the neural responses recorded from different subjects into a joint coding subspace^15,16^. This allowed us to jointly analyze all recorded trials and to use SVM decoders to accurately identify the subspace dimensions that contained avoidance-predictive (Fig. 3C) and tone-related activity (Fig. 3G). Furthermore, our sequential decomposition approach based on nullspace projections ensured that the identified dimensions were orthogonal, which isolated the different signals from each other (Fig. 3E).

While previous work already showed that mPFC activity contains avoidance-predictive information^8^, our approach allowed us to identify and characterize the activity patterns that carry this information. Jercog et al.^8^ reported that the prediction of avoidance actions was dependent on tone-responsive cells, suggesting joint coding of sensory and behavior-related signals. In contrast, our subspace decomposition revealed that avoidance-predictive activity was distinct from tone-related activity and that tone-related activity did not differentiate between avoidance and error trials (Fig. 4A). Thus, our results speak against a systematic overlap in the coding of sensory and behavior information. Instead, our results are consistent with a model in which sensory-driven mPFC responses partake in a distributed dynamical process^17^ to drive behavior execution. While our results are only correlational, multiple studies have demonstrated the causal role of mPFC in active avoidance^7,8,18,19^ via projections to the basolateral amygdala (BLA) and the nucleus accumbens^7,20^. A thorough understanding of how dynamic prefrontal computations carried out by distributed population signals map onto subpopulations projecting to different output regions will require further experimental and theoretical studies^21^.

Our result that activity in the tone dimension is modulated by behavior execution (Fig. 7) indicates that tone-driven mPFC signals are not purely sensory but are modulated by the behavior of the animal. The drop of tone-driven activity at action onset, despite continued sensory input, indicates a change of information flow induced by the execution of the learned avoidance action. However, it is unclear what causes the observed drop of activity. mPFC tone responses are dependent on inputs from the BLA^8^. Furthermore, the BLA is generally required for avoidance learning^22^ but is also involved in the expression of avoidance behavior^7,23^. The bidirectional interaction between BLA and mPFC offers a potential substrate for the observed dynamics of mPFC tone signals, but further work is needed to understand the temporal dynamics of sensory information flow and its contribution to behavior execution.

Finally, the switch between the two active avoidance actions (x- and y-shuttling) allowed us to reveal a neural substrate of behavioral flexibility in mPFC. mPFC has previously been shown to be involved in switching between tasks or rules^24–26^, and our results offer new insights into how behavior-related neural activity is updated upon a switch between conditioned behavioral responses. We found that avoidance-related activity was organized into two dimensions, where one was general to both avoidance behaviors, while the other was specific to y-shuttling and only emerged in task 2 (Fig. 5, Fig. 6). This sequential layering of previously learned transformations and the newly added dimensions might help animals not only to maintain the memory of previously learned tasks but also to shift between tasks in a context-dependent manner. In fact, the similar temporal dynamics of activity in motion and avoidance dimensions (Fig. 4A, B and S5) could indicate that with progressive learning new correlates of avoidance behavior are derived from either naive or previously learned behavioral primitives. The high level of mixed selectivity in mPFC should greatly facilitate such layered learning and future work should investigate how a context-specific recombination of sensory and behavioral neural correlates might facilitate behavioral flexibility.

## Methods

All animal procedures and experiments were approved by the Cantonal Veterinary Office Zurich, Switzerland.

### Subjects

All experiments were performed on male C57Bl6/Crl1 mice (Charles River Germany) aged between 4 to 7 months at the start of the behavioral experiment. Animals were housed in individually ventilated cages (IVC) in a 12 h light/dark cycle room (lights on from 7:00 to 7:00), and were provided food and water ad libitum. After import from the breeders, mice were given a two-week acclimatization period to the new housing condition prior to the first surgery. During the experiments mice were kept in groups of 2 to 5 animals.

### Surgical procedures

#### Anesthesia

For all procedures including anesthesia mice received pre-emptive buprenorphine (Bupaq; Streuli, 0.1 mg/Kg) 20-30 minutes prior to anesthesia. Anesthesia was induced with a Ketamin-Xylazin cocktail (Ketanarcon; Streuli, 90 mg/Kg / Xylazin; Streuli, 8 mg/Kg), and mice were mounted onto a stereotactic frame (Kopf Instruments). During the procedure mice received 95% medical O2 (Pangas, Conoxia) through a face mask and body temperature was kept steady at 37 degrees Celsius using a temperature controller and a heating pad.

#### Viral injections

At the time of the first surgery mice were 8 - 13 weeks old. To label excitatory neurons in the prelimbic cortex we intracranially injected 500nl (Titer: 4x10E11) of an adeno-associated virus driving the expression of GCaMP6m via the CamKII-promoter (AAV2/5-CamKIIa-GCaMP6m) into the prelimbic cortex (AP 1.8 ML 0.4 DV 2.1). We used either a micropump (UMP3UltraMicroPump; WPI) or a borosilicate glass pipette with a 50 μm diameter tip and injected the virus by applying short pressure pulses at a speed of approx. 100 nl/min. After injection the needle/glass pipette was left in place for 5 min to avoid backspill. Finally, the skin was closed using surgical sutures.

#### Microendoscope implantation

7 to 14 days after the viral injection we implanted a small stainless steel guide tube (1.2 mm diameter; Ziggy’s tubes and wires) with a custom glass coverslip (0.125 mm thick BK7 glass, Electron Microscopy Science) glued to one end as previously described in 27. In brief, we first made a 1.2 mm diameter (round) craniotomy centered above the ventral-medial prefrontal cortex (1.8 mm anterior, 0.4 mm medial, relative to bregma). To avoid increased intracranial pressure when inserting the implant, we aspirated tissue down to a depth of 1.9 mm from the skull-surface. Next, we lowered the guide tube to the bottom of the incision (2.2 mm relative to Skull surface) and glued the guide tube to the mouse skull using UV curable glue (4305 LC; Loctite). We then applied dental acrylic (Metabond; Parkell or Scotchbond ESPE; 3M) over the complete cranium and around the guide tube. Finally, we attached a metal bar and applied dental acrylic cement (Paladur) to stabilize the implant.

#### Analgesic regime

For 3 days after each surgical procedure animals received Buprenorphine s.c. (Bupaq; Streuli, 0.1 mg/Kg) every 6 h during the light cycle and in the drinking water (Bupaq; Streuli, 0.01mg/mL) during the dark cycle, as well as Carprofen s.c. (Rimadyl; Zoetis, 4mg/Kg) every 12 h.

#### Preparation of animals for behavioral experiments

Animals received 6-12 weeks recovery time before testing viral expression levels. Approximately one week before starting behavioral experiments we inserted the gradient index (GRIN) lens into the guide tube (GT-IFRL-100-101027-50-NC; GRIN-Tech) and attached a microscope base plate (Inscopix) above the implanted microendoscope with blue light curable glue (Flow-it; Pentron).

### Validation of imaging methodology

#### Perfusion

After completion of experiments animals were given terminal anesthesia with Pentobarbital (Esconarkon; Streuli, 200 mg/Kg) and perfused transcardially with PBS followed by 4% paraformaldehyde (PFA). Brain tissue was removed and post-fixed for 24-48 h in 4% PFA. Coronal slices (50 μm thick) were prepared on a Vibratome (VT1000 S; Leica) and stored in PBS.

#### Verification of microendoscopic implant

To confirm placement of the GRIN lenses in the medial-prefrontal cortex, cyto-structural differences in the tissue were highlighted using Nissl stain (NeuroTrace 530/615; Invitrogen) following the provided protocol from Invitrogen with a dilution of 1:50 NeuroTrace. Slices containing the prefrontal cortex were mounted and images were acquired using a fluorescence microscope (Olympus, BX51). Images were overlaid using the reference pictures from 28. For each section we marked the position of the base of the microendoscope for every mouse (see Fig. S2B).

#### Verification of cell type

Standard immunofluorescence protocols were used to stain inhibitory and excitatory neurons. Slides were incubated with the primary antibody (either rabbit anti-Neurogranin (1:2000, 07-425 Millipore) or rabbit anti-GAD65 (1:500, AB1511, Millipore)) at 4° C overnight followed by a 2-hour incubation at room temperature with the secondary antibody Alexa 594 anti-rabbit (1:200, A-11062 Invitorgen). Slides were further stained for 4 min with DAPI (1:1000, D1306, Invitrogen) in PB (0.1M) prior to mounting. Confocal pictures were taken in the red (at wavelength 594 nm, Neurogranin or GAD65), green (at wavelength 488 nm, GCaMP6m) and blue channels (at wavelength 390 nm, DAPI) and pictures were compared for overlap of labeling (see Fig. S2C, D).

### Behavioral procedures

#### Calcium imaging during mouse learning behavior

Calcium imaging experiments were performed using a miniaturized fluorescence microscope (nVista HD 2.0; Inscopix). Before behavioral experiments, we habituated all mice to the mounting procedure and the weight of the miniscope for at least three consecutive days. During the mounting procedure, animals were briefly head-fixed by fixing their metal head bar to a custom made mounting station with a running disc. Additionally, subjects were habituated to the experimental room and were handled by the experimenter for five out of seven days preceding the experiment. In every imaging session we verified for absence of shifts in the field-of-view and slightly adjusted the microscope focus if necessary. We acquired frames of 1000 × 1000 pixels at 12 bit and a frame rate of 20 Hz. To acquire the calcium imaging data, we used an LED intensity between 10 and 25% (100-150 μW) depending on the strength of the GCaMP6m expression. For all recordings we used the maximum imaging sensor gain level of 4. All recorded data was directly streamed to the hard-disc of a desktop computer.

#### Two dimensional active avoidance

For the two-dimensional active avoidance experiments we used a rectangular shuttle box (Cambridge Instruments) which we separated into four compartments by using four equally sized platforms. We 3D-printed these movable platforms to fit between the bars of the shock grid which allowed us to dynamically adjust the safe zone during training. In the default position, the platforms were situated below the shock grid such that mice could not jump onto them to avoid contact with the shocking grid. In the elevated state, mice had the possibility to fully stand on the platform without being in contact with the shock grid, thereby creating the possibility to avoid or escape shocks. We controlled the platforms using servo motors that we placed outside of the isolation chamber. The complete learning paradigm had a duration of 11 days comprising habituation (day 1), active avoidance task 1 (days 2-4), active avoidance task 2 (days 5-9) and extinction sessions (days 10 and 11). All sessions had a duration of 40 minutes and contained 50 trials with pseudorandom inter-trial intervals of 30±10 s. Each of the trials started with the presentation of an 8 kHz tone at 80 dB for 10 seconds. In all active avoidance sessions (days 2 to 9) the tone was followed by a light foot-shock (0.2 mA) with a maximal duration of 5 s. For each of the trials, we defined half of the shuttle box as a safe zone. We determined the position of the safe zone by the trial type (task 1 or task 2) and the position of the animal at trial start. For task 1 trials, mice had to cross the midline along the x-axis (x-shuttles) of the cage to reach the safe zone, whereas for task 2 trials, mice had to cross the midline along the y-axis (see Fig. 1A). If mice entered the safe zone during tone or shock presentation, we blocked both tone and shock channels until the end of the trial. If mice did not shut off the tone before shock onset, we elevated the two platforms in the safe zone for a duration of 15 s, time-locked to the onset of the shock, providing mice with the possibility to escape. We recorded all mouse behavior using two top view B/W cameras (DMK 23FV024; ImagingSource) which covered the entire cage and were later merged together to produce a single behavior video. The recording of individual frames of the behavior cameras was synchronized to the miniscope recordings using a hardware trigger, which allowed exact alignment of neural and behavioral data.

### Extracting neural activity from calcium imaging data

#### Preprocessing of calcium imaging data

We implemented the following procedures to preprocess the video of each individual imaging session. We first spatially down sampled all frames by a factor of 2 to obtain 500x500 pixel frames. Next, we used the TurboReg algorithm^29^ for motion correction by aligning each frame to a reference frame. We then temporally down sampled videos by a factor of 4, resulting in a frame rate of 5Hz. To account for slow changes in luminosity related to bleaching, we fit a rank-2 bleaching model by running PCA on a temporally smoothed version of the video and then subtracting this model from the original video. Next, to remove wide-field luminosity fluctuations occurring on a faster time scale (e.g. neuropil signals), we normalized each frame by dividing it by its lowpass-filtered version. Finally, we re-expressed all frames in units of relative changes in fluorescence, given by ΔF(t)/F0 = (F(t) – F0)/F0, where F0 is the mean frame obtained by averaging over the entire movie.

#### Cell extraction for individual sessions

To automatically identify individual neurons in the calcium imaging movies of a given imaging session, we used a well-established cell extraction algorithm based on principal component analysis and independent component analysis (PCA/ICA)^30^. This algorithm generates spatial filters that correspond to the cells’ locations, which allowed us to extract the corresponding temporal activity traces. However, instead of extracting these activity traces for each session individually, we first use the positional information contained in the identified spatial filters to align the movies from all imaging sessions of a given mouse.

#### Session alignment

To be able to track cells across imaging sessions we applied the following alignment procedure for each mouse. We first constructed cell maps for every session by calculating the maximum projection of all cells’ spatial filters onto one image (see outlines in Fig. 1F). We then used MATLAB’s *imregister* function to align the sessions’ cell maps onto one reference session. Next, we used the resulting registration coordinates to align all session movies into a common reference frame. This allowed us to concatenate all session movies to construct one movie containing the full experiment. To account for differences in the signal to noise ratio of individual sessions, we calculated the overall standard deviation of all pixels for every session and then scaled the corresponding movies to match the minimal standard deviation. The resulting concatenated movie thus contained ΔF/F values with a stable mean and standard deviation over all sessions.

#### Joint analysis of multiple sessions

We used the aligned and concatenated movies of individual subjects and PCA/ICA to obtain spatial filters and activity traces over the whole experiment. Since the high number of frames made running PCA/ICA on the whole concatenated movie intractable, we instead generated spatial filters by performing signal extraction on a reduced movie, containing 6000 consecutive frames from every session (i.e. half of the data). We then recovered the activity traces over the full duration of the concatenated movie, by projecting the full movie onto these spatial filters.

#### Post-processing and validation

A known issue with PCA/ICA is that individual cells are occasionally split into multiple components. To make sure we do not include split cells in our analyses, we detected pairs of cells that have highly correlated activity (Pearson correlation > 0.7) and are spatially close (centroid distance < 20 pixels) and excluded one of the cells for each pair. Finally, we manually validated each cell by inspecting its morphology, activity trace over all sessions, mean calcium transient, and by checking whether peaks in the activity trace were consistently caused by the same pixel pattern.

### Quantification and statistical analysis

#### Behavior analysis

To analyze animal behavior, we first stitched the videos of the two behavior cameras to obtain a single video. We then used the DeepLabCut software^31^ to track five points of the animal (see Fig. S1). To quantify the overall speed of the animal, we averaged the positions of the three most stable points (left ear, right ear and miniscope bottom) and calculated the instantaneous speed per time step.

#### Alignment of trials and ITI shuttles

We aligned avoid trials according to the start of the avoidance action (avoid start). We defined the action start time as the time point with the maximal increase in instantaneous speed within the 2 s window before the detected shuttle. For all analyses that considered the window starting 3 s before action start, we discarded avoidance trials with an action start earlier than 3 s after tone start. To ensure that error trials were comparable to avoid trials in terms of trial lengths, we randomly sampled error trial alignment points, such that they matched the distribution of avoid start time points between 3 s and 9 s. To account for the variability introduced by this sampling, we repeated each analysis over multiple repetitions (see decoding). We aligned ITI shuttles analogously to avoid trials.

#### Single cell analysis

To define cells as trial-responsive, we considered the 3 s window after tone start and the 3 s window before action start. We calculated z-scores per time step, using the 6 s window before trial start as a baseline period. We defined cells as trial-responsive if their mean absolute z-score exceeded a value of 1.96 (p < 0.05, two-tailed)^8^.

#### Subject alignment

To align the population activity of different subjects into one common subspace we first collected event-aligned trial averages (see Fig. S3). We separately aligned data from avoid trials, error trials and the ITI for the two tasks (i.e. 2x3 conditions). For avoid trials we used windows around the tone start (-1 s to 3 s) and action start (-3 s to 1 s) alignment points. For error trials we used the same structure using the sampled alignment points (pseudo-action start). For ITI shuttles we used the window from -4 s to 4 s around action start. We next computed condition averages for each cell in each of the six conditions and concatenated all cells from all animals to obtain six (n x t) matrices, where n is the total number of neurons and t the number of time steps (8 s at 5 Hz). We then mean-subtracted these six matrices. To prevent avoidance-related and motion-related activity from being mapped into the same dimensions solely due to their temporal overlap in avoidance trials, we performed an orthogonalization step to eliminate motion-related activity in the avoid condition averages. We performed the following procedure individually for the condition averages from the two tasks. First, we performed PCA on the ITI condition averages. Next, we calculated the nullspace of the resulting PCs. We then projected the avoid condition averages into this nullspace to remove motion related information. Finally, we concatenated the two resulting corrected avoid condition averages with the remaining 4 condition averages along the time dimension to obtain an (n x 6t) matrix on which we then performed PCA. We defined the resulting (n x k) matrix of coefficient values as the joint subspace, where k is the number of PCs we choose to use. To compute subject-specific projection matrices into this joint subspace, we split the coefficient matrix back into coefficient matrices for the individual subjects. Since these matrices are not orthogonal anymore, we used the QR-decomposition to orthogonalize them as the final step of the procedure. To ensure that the alignment procedure did not introduce artifacts in further analyses we used half of the trials for alignment, and the other half for the decoding analyses described below. The choice of trials was randomly assigned for every repetition.

#### Avoid/error decoding

We aligned avoid and error trials to (pseudo) action start and trained individual decoders for every time step from -3 s to 1 s from the alignment points. Decoders were linear support vector machines and we used 5-fold cross-validation to estimate test-accuracies. We used avoid and error trials from days 3 to 9 and balanced the two classes by subsampling 300 trials per class in all settings. To deal with the variability introduced through sampling (error trial alignment and trial samples), we repeated each analysis multiple times (typically 50 times, if not reported otherwise) and computed average accuracies over repetitions.

#### ITI control and video decoders

In the ITI decoding setting we considered the window from -3 s to 1 s around avoid start for each ITI shuttle and trained decoders to discriminate them from random 4 s periods in the ITI. For the video decoder control setting we used the 5-dimensional speed vector from the DeepLabCut tracking points instead of the 10-dimensional neural activity vector. In order for the ITI decoding setting to be a useful control, we needed to choose the ITI data such that the decoding performance of video decoders are comparable between the ITI and the avoid versus error settings. By matching the performance for the video decoders, we could interpret potential differences when using the same data for the neural decoders. To match the performance of video decoders for the two settings, we sorted ITI shuttles according to their speed (fast shuttles are easier to decode) and chose the 11 fastest shuttles for every session. This choice resulted in the performance reported in Fig. 2.

#### Identification of motion dimensions

To identify motion-related dimensions in the joint coding subspace we collected ITI shuttles from task 1 and task 2 sessions and computed average activities for the window from -4 s to 4 s around avoid start. We then performed PCA on the resulting activity matrix and considered the first 5 PCs as motion dimensions.

#### Identification of avoidance dimensions

To define avoid dimensions we first projected all trial data into the nullspace of the first two motion dimensions. Next, we iteratively defined avoid dimensions using the following procedure. 1) We train a time-independent avoid/error decoder using randomly sampled time points from the 4 s window around action start (1 per trial). 2) We compute the avoid dimension by normalizing the decoder weight vector to have a norm of 1 and projecting this vector back from the nullspace into the 10-dimensional subspace. 3) We project the trial data into the nullspace the space given by the first two motion dimensions and all avoid dimensions. 4) We repeat the process with different trial samples and time-point samples until we have obtained five avoid dimensions.

#### Identification of tone dimension

To identify tone dimensions we followed the same strategy as for avoid dimensions in the nullspace of the first two motion dimensions and the first two avoid dimensions. We trained a time-independent tone decoder using randomly sampled time points during avoid (before action start) and error trials. The baseline period was defined using data points from the period 1 s before tone start.

#### Task decoding

We trained time-dependent SVM decoders to discriminate between data from task 1 (days 3 and 4) and task 2 (days 6 to 9) based on activity in the 5-dimensional coding space. We trained an independent set of decoders for avoid trials, error trials and ITI shuttles. To quantify the importance of a given coding dimension for task-decoding accuracy, we projected trials into the nullspace of this dimension, repeated the decoding procedure and quantified the change in accuracy.

### Software and Statistical Tests

For all data analysis (image pre-processing and population analysis) and statistics we used the MATLAB programing environment. If not indicated differently we used non-parametric statistical tests. Box plots indicate median (center), 25^th^ and 75^th^ percentile (box) and most extreme data points (whiskers) that were not considered outliers (points for which the distance from the box exceeds 1.5 times the length of the box). Violin plots indicate mean and standard deviation. The sample sizes required for this study were initially estimated based on pilot behavior studies, but no formal statistical tests were used to predetermine sample size. We excluded one animal because we did not observe any neuronal activity, due to insufficient labeling and/or GRIN lens misplacement. We had to terminate the behavior experiments for 6 mice to be in accordance the animal welfare regulations because they did not learn the task sufficiently (performance below 50% after 3 days of training). We excluded 4 imaging sessions (from a total of 132, 12 mice x 11 days) because we could not align the recorded frames to frames from previous sessions. All excluded sessions were from the extinction phase at the end of the experiment (days 10 and 11).

## Acknowledgements

We would like to thank Elisabeth Abs, Reinhard Loidl and Pau Aceituno for fruitful discussions and for feedback on the design of the study and the interpretation of the results. We would like to thank Simone Holler-Rickauer for the preparation of the histology slides and Jasmine Kofmel and Lou-Ann Ingold for help with the verification of cell labeling.

## Author Contributions

B.E. and B.F.G. conceptualized the study, B.E. carried out all in vivo imaging experiments and data analyses, C.H. helped building the behavior setup, M.C., V.M. and E.A. provided conceptual feedback for behavior experiments and population analysis, R.B. helped with the behavior experiments, animal preparation and DLC tracking, V.M and A.G. conceptualized the subject alignment procedure, B.E., E.A., R.B. and B.F.G. wrote the manuscript.

## Supplementary Figures

**Figure S1:**
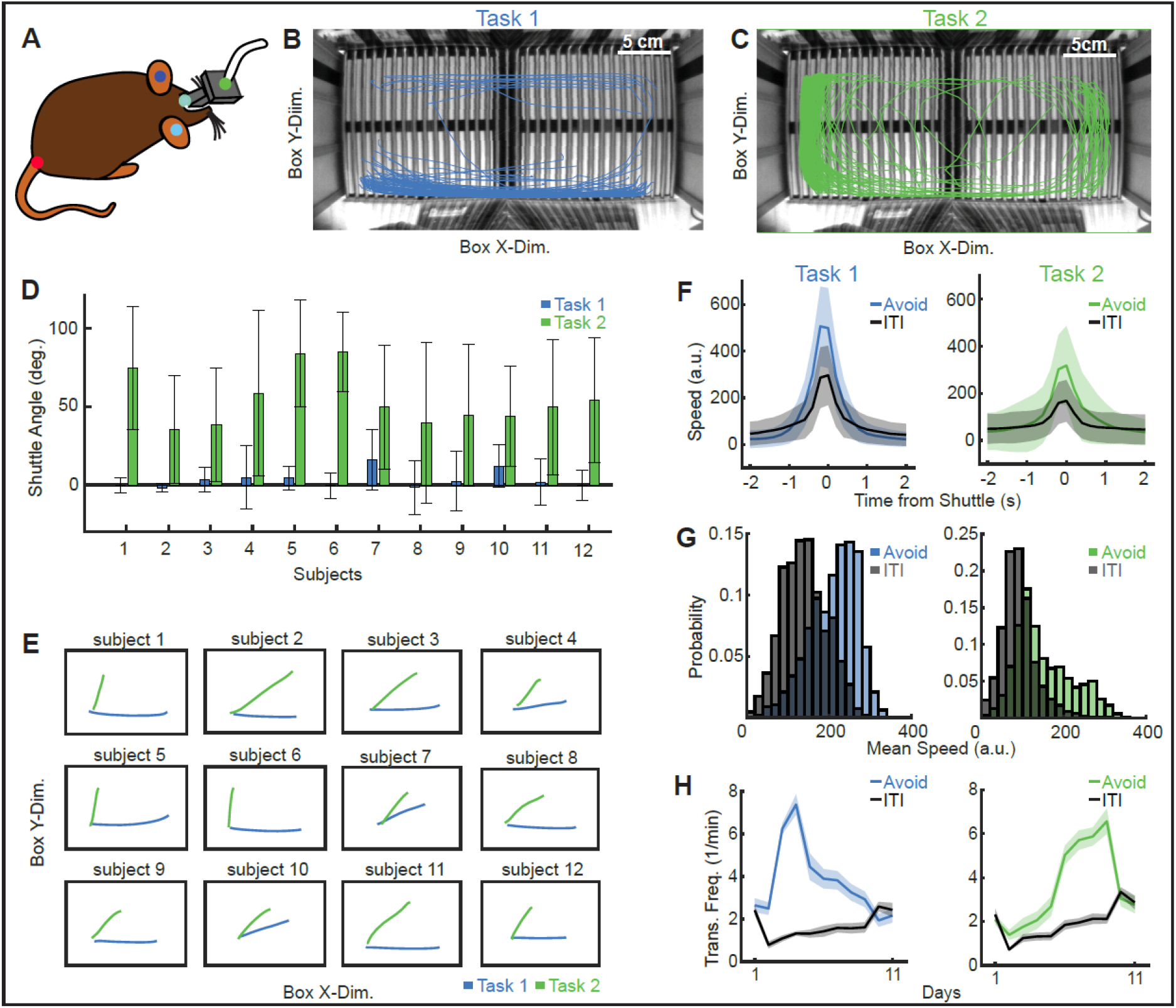
Animal tracking and behavior analysis. (**A**) DeepLabCut tracking points of the animal. (**B, C**) Avoidance trajectories during task 1 (x-shuttling) and task 2 (y-shuttling), respectively. (**D**) Distribution of shuttle angles in degrees for all 12 animal subjects in tasks 1 and 2 (mean ± s.e.m over avoid trials). (**E**) Mean shuttle trajectories depicted in the shuttle box for all 12 animals. (**F**) Distributions of mean transition speed for ITI (gray) and avoid (colored) shuttles in tasks 1 (left, 1151 avoid shuttles, 1329 ITI shuttles) and 2 (right, 1945 avoid shuttles, 3859 ITI shuttles). (**G**) Animal speed time-courses aligned to the shuttle start for tasks 1 (blue, left) and 2 (green, right) (mean ± s.d. over shuttles from 12 mice). ITI shuttles are depicted in gray. (**H**) X-axis (blue) and Y-axis (green) shuttle transition frequency compared to the respective ITI (gray) shuttle frequency across days (mean ± s.e.m. over 12 mice).

**Figure S2:**
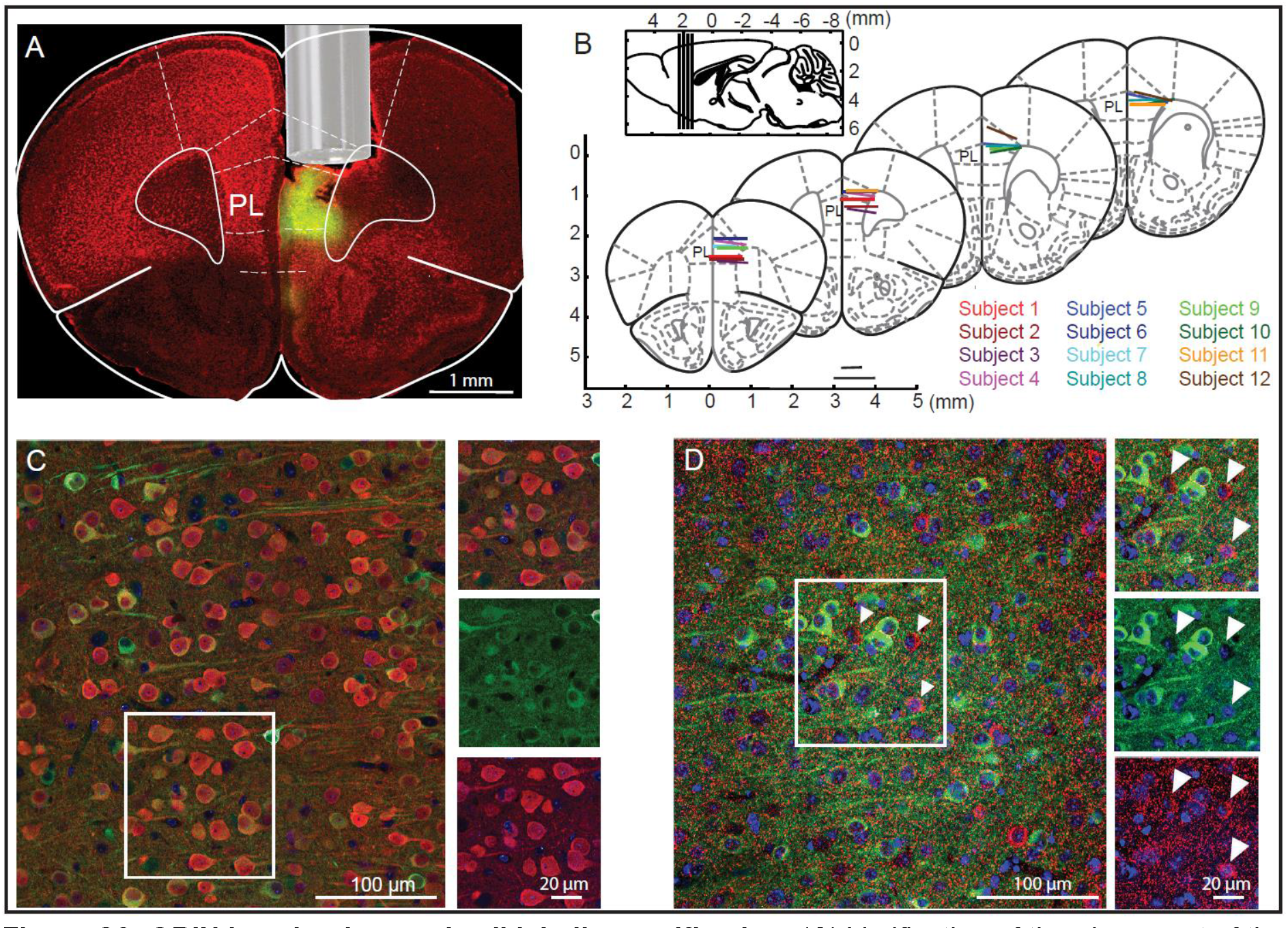
GRIN lens implant and cell labeling verification. (**A**) Verification of the placement of the GRIN lens, with the GRIN lens position displayed in an example coronal mouse brain section. Nissl stain (red) and GCaMP6m (green). (**B**) Locations of the GRIN lens surface mapped onto the mouse brain atlas for each individual animal. (**C**) Left: Immunohistological validation of GCaMP6m expression in PL, comparing GCaMP6m (green) and neurogranin labeling (red) in excitatory neurons. Top right: overlap. Middle right: GCaMP6m. Bottom right: neurogranin. (**D**) Left: Comparison of GCaMP6m expression (green) to GAD65 staining (red). Top right: overlap. Middle right: GCaMP6m-expressing neurons. Bottom right: GAD65-positive neurons. Arrows indicate three example GAD65-positive neurons.

**Figure S3:**
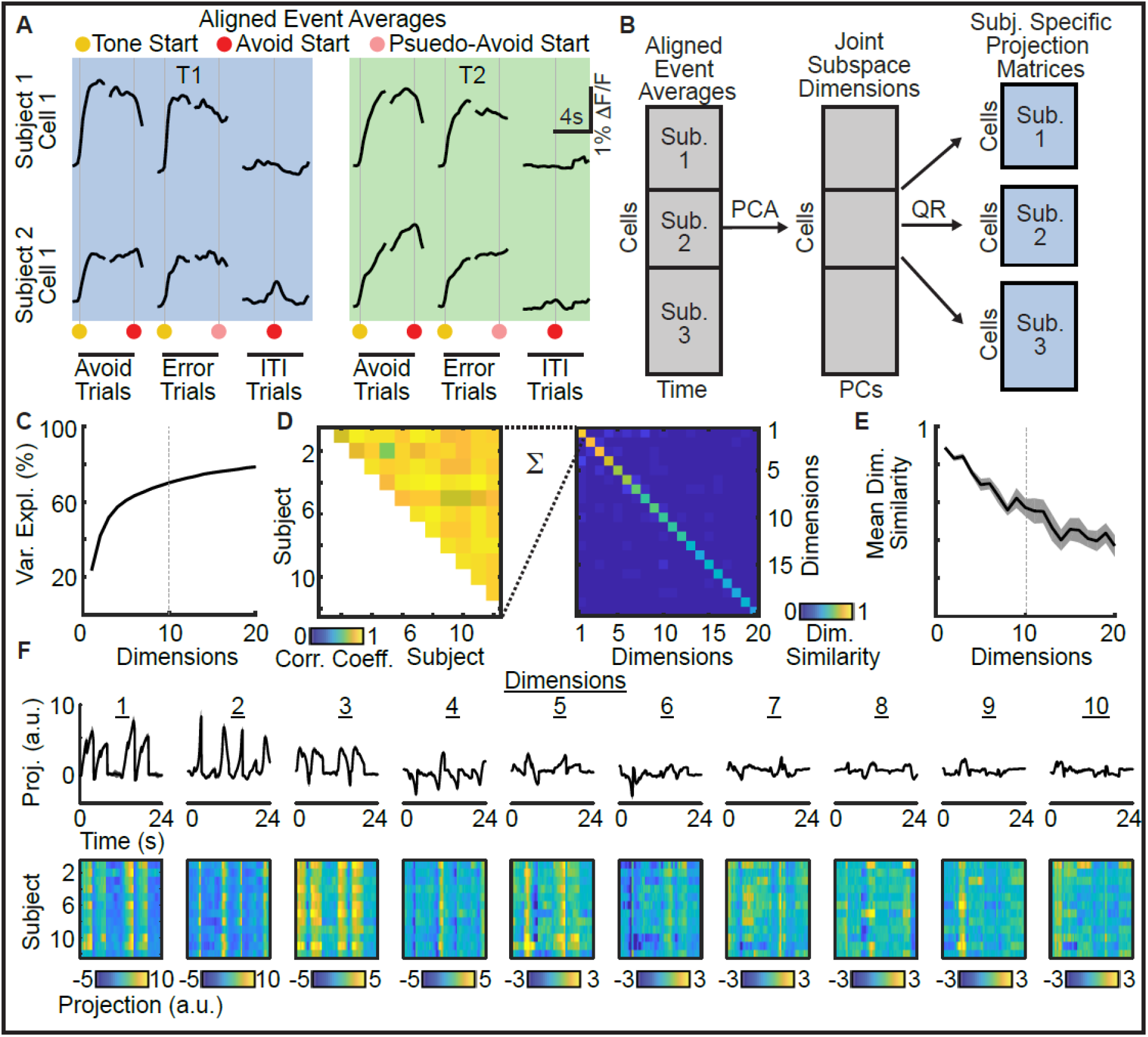
Alignment of neural data across animals into a joint subspace. (**A**) Specification of time points used for alignment, displayed for two example neurons from different subjects that show similar responses during avoid, error and ITI events in task 1 (blue shade) and task 2 (green shade). (**B**) To align neural population data, the temporally aligned event averages displayed in A) are first concatenated for all cells. We then process these event averages (see methods) and concatenate them along the time axis. Next, PCA is used to generate the joint subspace which is defined by the coefficients of the first n PCs (n is chosen below). Subject-specific projection matrices into the joint subspace can then be computed by splitting the coefficient matrix back into matrices for individual subjects and orthogonalizing them using the QR decomposition. (**C**) Variance explained by the first 20 subspace dimensions (mean ± s.d. over 50 repetitions). In this work, we include only the first 10 PCs into the joint subspace. (**D**) Left: Cross-subject correlation of the first subspace dimension. Right: Similarity of pairs of dimensions, where similarity is computed by averaging the elements of the triangular cross-subject correlation matrix displayed on the left. (**E**) Average dimension similarity for the first 20 subspace dimensions (mean ± s.d. over 50 repetitions). We chose the number n of used dimensions to be 10, as it constitutes a good tradeoff between explained variance (see C) and alignment quality. (**F**) Top: Mean projections onto subspace dimensions 1 to 10 (n = 12 subjects). Bottom: Projections displayed for individual subjects, highlighting common structure.

**Figure S4:**
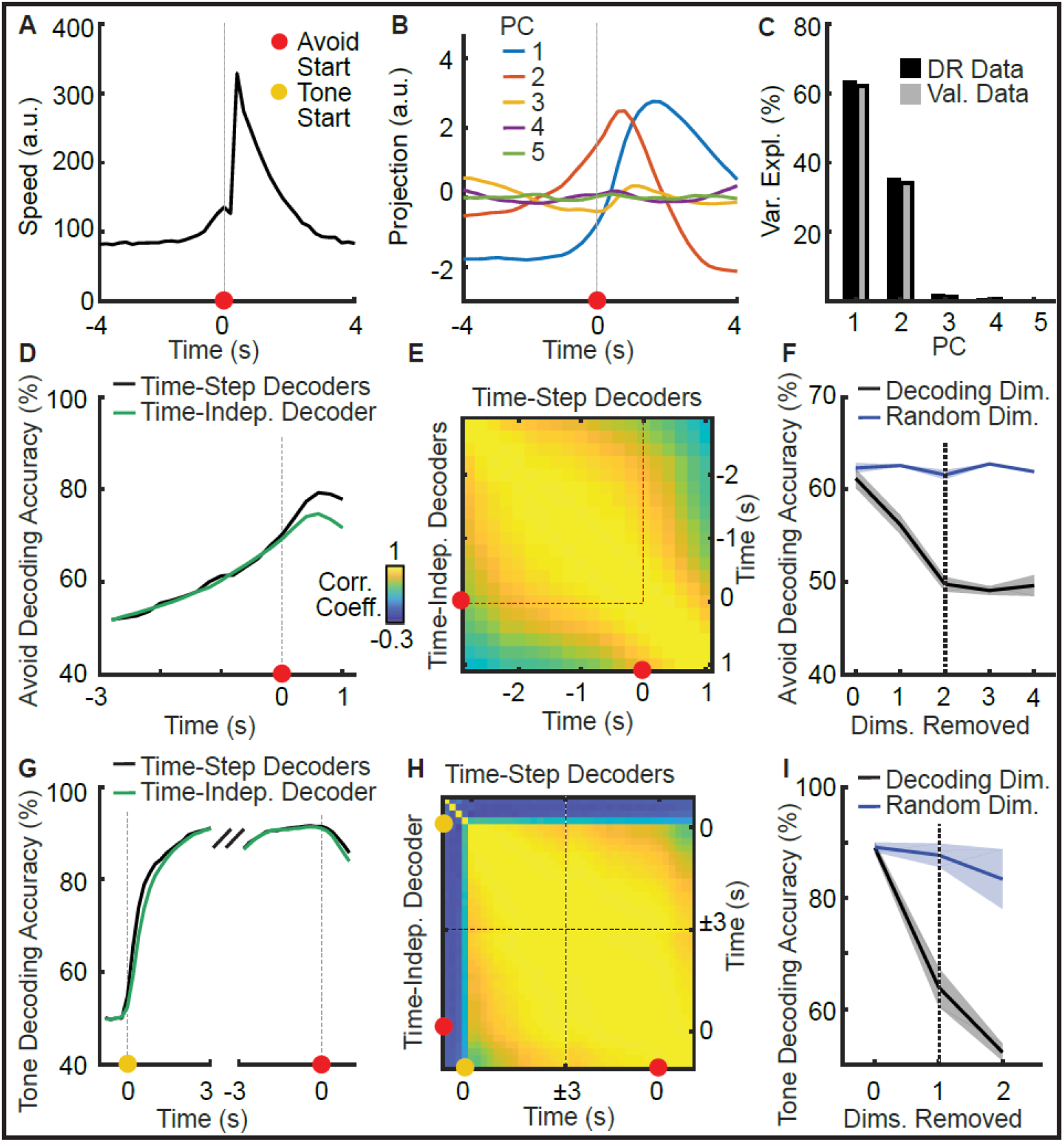
Definition of coding dimensions. (**A**) Mean animal speed during ITI shuttles used for ITI shuttle decoding (n=1452 ITI shuttles). (**B**) Principal components of neural activity in the joint subspace averaged over ITI shuttles used for the definition of motion dimensions. (**C**) Variance explained by the first five PCs for trials used in dimensionality reduction (DR data) and held-out validation trials (Val. data). (**D**) Accuracy of avoid versus error decoding per time-step for a set of time-dependent decoders trained individually per time step, and one single decoder trained using data from all time-steps (mean over 50 repetitions). The single decoder was separately evaluated with data from different time steps. The resulting accuracies match the performance of individual decoders and becomes worse only after action onset, suggesting stable avoidance predictive activity patterns prior to action onset. (**E**) Correlation coefficients for pairwise comparisons of time-step decoder weights (mean over 50 repetitions). Especially before action onset decoder weights show high correlations, indicating a stable representation of avoidance-predictive activity. (**F**) Decoding accuracy of time-independent decoders for the progressive removal of avoid dimensions (mean over 50 repetitions). Avoid dimensions are defined by the decoder’s weights vector and are iteratively removed via nullspace projections. Comparison to randomly removed dimensions in blue. (**G-I**) Analogous to (D-F) for tone decoding. Time-step tone decoders show representational stability and performance of time-step decoders can be matched using a single time-independent decoder.

**Figure S5.**
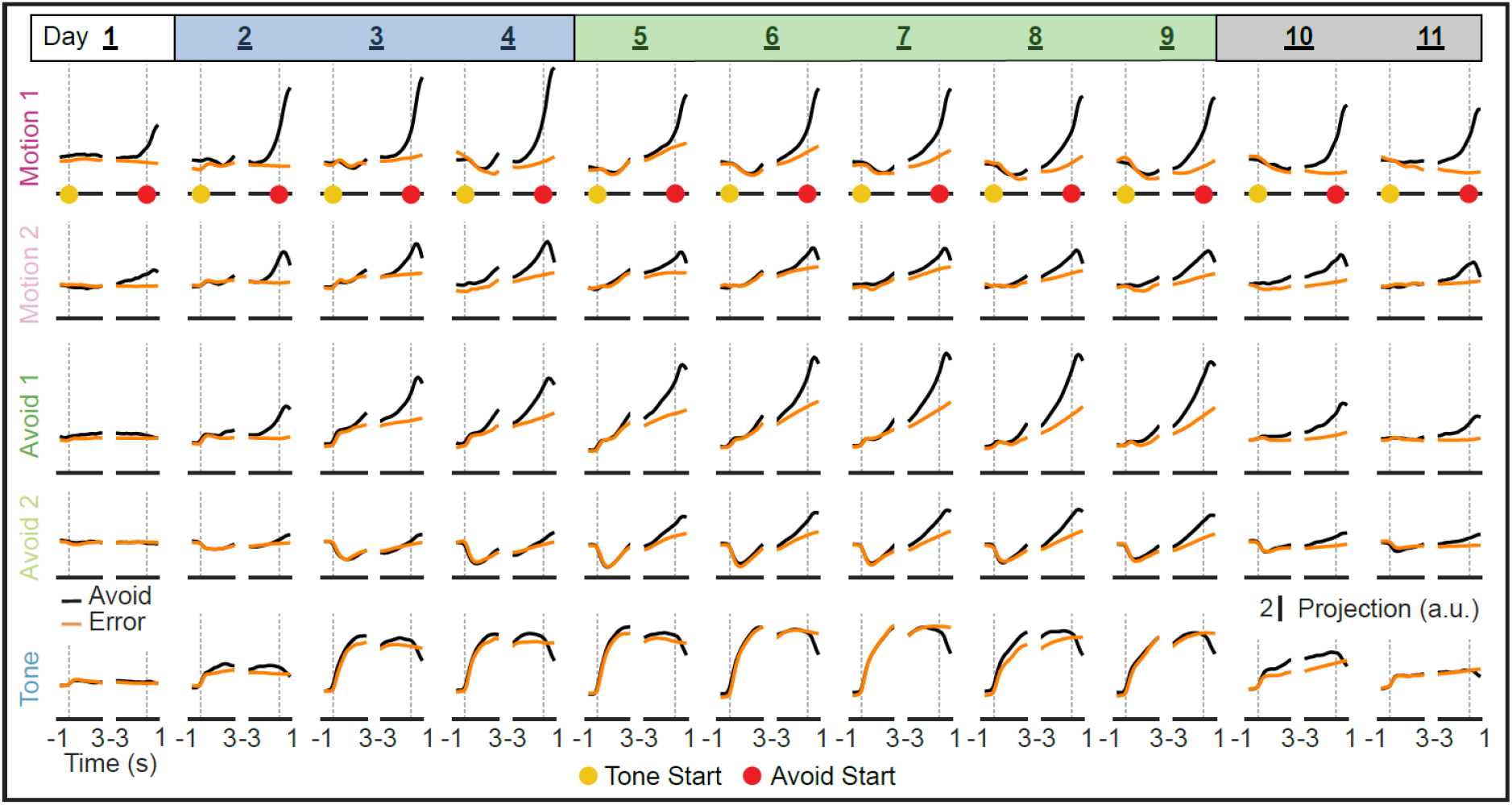
Coding dimension projections per session. Mean projection (n = 50 repetitions) of the five coding dimensions (as presented in Fig. 4) displayed across all 11 days of learning for avoid (black line) and error (orange line) trials.

